# Cascading carry-over effects of early activity phenotypes in golden eagles

**DOI:** 10.1101/2025.09.21.677571

**Authors:** Svea-Sophie Zimmermann, Martin U. Grüebler, Julia S. Hatzl, Enrico Bassi, Kamran Safi, Wolfgang Fiedler, David Jenny, Matthias Tschumi

## Abstract

Early-life conditions are often associated with fitness of animals later in life. Although recent studies show that environmental conditions during development affect the formation of early behavioural phenotypes we have a limited understanding of the cascade of life-history transitions from early phenotypes to fitness relevant behaviours in later life stages. Here, we used GPS and body-acceleration data of 35 juvenile golden eagles (*Aquila chrysaetos*) to investigate the association between nestling body condition and activity levels (ODBA), the relationship of nestling activity with the timing of fledging and post-fledging activity, and the effects of post-fledging activity on post-fledging movements and the timing of dispersal. We found that nestlings with good body condition also showed increased activity levels. Active nestlings fledged earlier and remained more active after fledging than passive nestlings during early post-fledging. Increased post-fledging activity levels were related to an increased number of pre-dispersal forays and an early timing of dispersal. Even though some effects showed reduced certainty, our results provide evidence for downstream effects of behavioural differences due to early-life conditions. They suggest that early activity phenotypes of nestlings drive the timing of subsequent life-history transitions, thereby representing a key mechanism that links early life conditions with future performance.

## 2. Introduction

During ontogeny, most organisms transition between life stages during which they are confronted with often fundamentally different environments [1]. Because of the challenges associated with new environments, early-life stages are characterised by high mortality, typically occurring just after transitioning from one life stage to another [2]. Often, survival is strongly affected by the early-life conditions experienced [3,4]. Early-life conditions can however also drive the expression and development of individual traits that subsequently carry-over to later life stages [5]. Such carry-over effects of early-life conditions can have crucial consequences on stage-specific individual performance affecting population composition and persistence [6,7]. In the last decades, research has provided fundamental insights into the prevalent associations between early-life conditions and fitness parameters later in life [5,8]. However, such within-individual associations often considered only two points, one at the beginning and one at the end of the entire ontogenetic trajectory, ignoring the translation processes of early life conditions over multiple life stages (but see [2,9]).

The mechanisms underlying carry-over effects of early-life conditions can be linked to physiological characteristics such as oxidative stress, corticosterone levels or telomere length [10,11]. However, early-life conditions can also shape behavioural phenotypes i.e. individual behavioural characteristics which then may be associated with future fitness [12]. Thereby, we are becoming increasingly aware of natal effects that affect future locomotor and cognitive abilities [13], but also social traits [14]. Yet, we still have limited understanding of the downstream effects of early behavioural differences over multiple life stages.

In altricial birds, fledging from the nest is one of the most fundamental life-history transitions, as it marks a crucial first step towards independence from the parents and initiates flight trough an unknown environment [15]. Growth, physiology and behaviours related to flight start to develop in the nest and developmental thresholds need to be reached for successful fledging [16,17]. Thus, the conditions experienced in the nest were found to affect the timing of fledging [18]. Recent findings suggest that conditions in the nest not only shape body condition and timing of fledging, but also impact behavioural phenotypes [14]. It therefore appears likely that behavioural differences associated with body condition are responsible for the translation of conditions in the nest to individual differences in the post-fledging period.

For territorial species with extended parental care, becoming independent and leaving the parental home-range after the post-fledging period (i.e. the start of natal dispersal) represents another fundamental life-history transition [19]. In many species, an early start to dispersal can facilitate its outcome by enabling the early detection and monopolisation of resources, floater areas and breeding sites [19]. Yet, individuals need to reach behavioural maturity to be successful [20].

Individual development is often not terminated at fledging but continues into the post-fledging period [16,17,21]. Beyond morphological and physiological developments, birds need to improve behavioural skills such as locomotion or vigilance [17,22]. As in the nestling period, developmental trajectories during the post-fledging period are expected to affect the timing of subsequent life-history transitions such as the departure to dispersal [23]. The post-fledging development is likely be shaped by both behavioural differences carried over from the nestling period and conditions experienced during the post-fledging period. While our understanding about the environmental factors affecting the timing of dispersal is increasing [24,25], the effects of behavioural differences on dispersal timing are not well established. This restricts our mechanistical understanding of early life effects on individual fitness components later in life and possible consequences for populations.

Behavioural differences are difficult to record for free-roaming species during the post-fledging period. However, animal borne accelerometers now allow quantifying individual behaviours. For example, overall dynamic body acceleration (ODBA) [26] has been shown to be a reliable proxy for activity-related energy expenditure [27]. ODBA can thus be used to quantify individual differences and developmental trajectories of time investment into energy-demanding behaviours such as locomotor training in the nest or flight behaviours after fledging [9,27]. Thereby, accelerometers allow investigating carry-over effects of early-life activity levels to activity levels in subsequent life stages [9] and enable novel mechanistic insights into how early life differences shape survival, recruitment success and distribution of behavioural phenotypes after independence..

Here, we investigate individual activity levels of nestlings as well as their carry-over effects over two major life-history transitions in Alpine golden eagles (*Aquila chrysaetos*), while accounting for sex differences and the environment of the parental territory. The Alpine golden eagle is a large territorial, non-migratory raptor which breeds in high-mountainous habitats [28]. Golden eagles largely rely on environmental energy for energy-efficient soaring flight [29]. After fledging, golden eagles show an extensive post-fledging period where flight behaviours mature [21,30,31] with large individual variation in the timing of permanent departure from the parental territories [22,23]. First, we quantified the correlation between individual activity levels in the nest and nestling body condition, suggesting that high activity levels reflect favourable nestling conditions. We also expected that high activity levels result in early timing of fledging. Second, we quantified the correlation between nestling activity levels and post-fledging activity levels, expecting that high activity levels in the nest translate into high activity levels after fledging. Finally, assuming that post-fledging activity levels reflect the development of flight behaviours, we analysed the effect of post-fledging activity on post-fledging exploration behaviour and the timing of departure. With this study we provide novel insights into the behavioural downstream effects of early-life conditions over two life-history transitions, with potentially far-reaching consequences for natal dispersal and recruitment.

## 3. Materials and Methods

### Study species

The Alpine golden eagle (*Aquila chrysaetos*) is a long-lived, territorial apex predator. With an average territory size of 50 km², they usually rely on multiple nests on cliffs or less commonly on trees, which are used alternatingly for many years [28,29]. Generally, pairs raise one to two chicks per brood, but breeding frequency fluctuates among years and pairs [28]. Chicks fledge at an age of 60 to 85 days between mid July and mid August and remain dependent on parental care for multiple months thereafter [22,28,29,32]. During this time they are mainly raised on fresh meat of Alpine marmot (*Marmota marmota*), chamois (*Rupricapra rupricapra*), ibex (*Capra* ibex), roe deer (*Capreolus capreolus*), mountain hare (*Lepus timidus*), and grouse species (Tetraoninae) but increasingly feed on carcasses of mostly ungulates [28,32]. After departure from their parental territory, golden eagles spend multiple years in transience before trying to take over or establish a territory [28,29,33].

### Study area

The study was carried out in the cross-national Alpine region covering eastern parts of Switzerland (Grisons) and north-eastern parts of Italy (Lombardy and South Tirol). The study area is characterised by a diverse topography of deep valleys and high mountain peaks with steep cliffs [28]. Forests often stretch along valley slopes up to elevations between 1’900 m and 2’300 m where they give way to open ground and vegetation dominated by grasses and heather. Golden eagle territories almost completely cover the Alpine range reaching high breeding densities [28].

### Tagging and tracking

Between 2017 and 2020, we equipped 35 juvenile golden eagles (2017: 2, 2018: 8, 2019: 18, 2020: 7) from 27 different territories with solar GSM-GPS-ACC loggers (e-obs Gmbh, Munich, Germany), using a leg-loop harness made of two-layered Teflon ribbon with a silicon core (to allow for relative elasticity of 20%). Tagging took place between late June and mid-July when nestlings were 52.3 ± 4.6 sd days old. The age of each nestling was calculated based on the hatching date that was assessed during brood monitoring. The combined weight of transmitters and harnesses of 70 g made up less than 3% of the birds’ body mass. In five territories, two siblings were tagged in the same nest; in two territories, single nestlings were tagged in the same nest but in two consecutive years and in one territory, single nestlings were tagged two years apart and in two different nests. In addition to tagging, we ringed, weighed and measured all the nestlings and collected feathers with blood quills for genetic sex identification.

Transmitters recorded GPS and tri-axial accelerometer (ACC) data from 04:00 to 17:00 UTC in summer (May-October) and 09:00 to 16:00 UTC in winter (November-April). GPS locations were recorded in 20-minute intervals. ACC data was recorded every five minutes in 7.9 second bursts, on three orthogonal axes resembling body planes (x, y, and z) at a frequency of 33.3 Hz (1188 bytes of data per burst). All data recorded by the tags were transmitted to and stored on the movebank data base [34]. For further analyses, GPS data were filtered to retain only the first location for every hour.

### Fledging, exploration and departure to dispersal

We defined the timing of fledging visually by identifying the first trajectory of consecutive GPS locations from the scattered GPS signal inaccuracies around the nest. We thus defined fledging time as the first timestamp of the trajectory leading away from the nest site. The age of fledging was then calculated by subtracting the hatching date from the date of fledging. To determine explorations and the onset of natal dispersal, we used a spatiotemporal threshold adapted from “Method 7” described in [35]: First, we calculated centroids from a kernel calculated by the R-package ‘*spatstat*’ [36], based on post-fledging locations clearly before emigration. Around these centroids, we selected an 8 km radius as a distance threshold to fully include the juveniles’ main home-range. The area within that threshold encompassed the juvenile home-range, favouring overestimation of smaller home-ranges to underestimation of large ones. Forays during the entire post-fledging period were defined as movements outside the circular home-range (minimum of one location) for a maximum duration of up to 14 days. The start of each foray was marked by the first location beyond the 8 km threshold, and the end point as the first subsequent location within the threshold. Adjustments were made based on visual examination of GPS tracks when a bird returned within the vicinity of > 6 km but < 8 km for no longer than two hours. In such a case, forays were counted as one instead of two. For each foray we calculated the duration in days and the maximum linear distance from the juvenile home-range centre in km. For the following statistical analyses, the number of forays was summed and the average duration of all forays, as well as the mean of all maximum distances calculated per individual during the entire post-fledging period. Departure to dispersal was defined as any movement outside the 8 km threshold without any overnight returns for more than 14 days (permanent departure) and the age at departure as the departure date minus the hatching date. Three of the 35 individuals were excluded from the foray and departure analysis as their tags either stopped transmitting (n = 2) or they left their parental territory only a few weeks after fledging (n = 1) without undertaking any forays.

### Juvenile activity

We used overall dynamic body acceleration (ODBA) as a measure of activity [26]. We transformed the raw ACC data into gravitational force *g* and calculated mean ODBA for each 7.9 second long ACC burst using the package ‘moveACC’ [37]. We determined ODBA during the nestling period for the 11 days prior to fledging and for the first 45 days of the post-fledging period. The 45 days after fledging correspond to the part of the post-fledging period before individuals start undertaking regular and longer forays and match with the duration of linear activity development [22,23]. We calculated daily mean ODBA including recordings from 06:00-17:00 UTC. Days with data gaps of more than one full hour (n = 77 days from 12 individuals) and days with early forays undertaken within the first 45 days after fledging (n = 37 days from 14 individuals) were excluded from following analyses. For the nestling period (11 days before fledging) and for the early post-fledging period (45 days after fledging) we fitted linear mixed-effects models to investigate the effect of sex and the relative number of days before and after fledging, respectively, on mean daily ODBA (Table S1). The predictor “day” (i.e. number of days before or after fledging) was centred and scaled for each period and included as a fixed effect as well as the random slope for each individual. We used the sex-corrected random intercepts derived from these models as indicator for individual ODBA during each period (Table S1; Figure S1). Additionally, random slopes during the post-fledging period indicated the individual development of daily ODBA over time (Table S1; Figure S1). By standardising the predictor “day” for each period, the random intercepts represented the average ODBA for each period and individual bird.

### Intrinsic and habitat variables

As a proxy for individual body condition, we calculated the deviation of recorded body mass from the mean body mass of males and females, respectively, to account for sexual dimorphism occurring in golden eagles [38]. We found no significant effect of age at tagging on body mass (LMM: Estimate = 45.65; 95% CrI = -108.07 to 190.42) and the age of tagging was thus not accounted for to calculate body condition. Nest exposure was measured as compass degrees and translated into northness (degrees) and eastness (degrees) for the analyses. To calculate a proxy for parental habitat quality, we used yearly mean values of the normalized difference vegetation index (NDVI) per juvenile home range as an indicator for food availability via trophic cascades [39]. To avoid overestimation of used space and foraging grounds, we calculated juvenile home ranges as 90% Minimum Convex Polygons (MCPs) using GPS points within the 8 km threshold defined above. During the post-fledging period, juvenile and parental home ranges are highly similar [40]. As Golden eagles mostly forage in open areas which reflect their main prey’s preferred habitat [28], closely forested areas were excluded (Forest Type 10 m High Resolution Layer derived from the Copernicus Land Monitoring Service). We extracted NDVI images using MODIS (Moderate Resolution Imaging Spectroradiometer) vegetation indices of the study area taken every 16 days from beginning of May 2017 to the end of April 2021 at 250 m resolution [41] and cropped them to the extent of each juvenile home range in QGIS 3.18 [42]. We then used the cropped NDVI images to calculate yearly mean NDVI values for each pixel (250 m x 250 m) which we subsequently averaged over the entire home range. Negative NDVI values representing unvegetated areas were set to 0.

### Statistical analysis

All statistical analyses were performed using R software version 4.0.5 [43]. We fitted Bayesian linear mixed-effects models (LMM) and generalized linear mixed-effects models (GLMM) using package ‘*rstanarm*’ [44]. To model nestling activity, we fitted an LMM for ODBA (random intercept of the ODBA model described above) during the nestling period as response variable, and nest exposure, NDVI, body condition and sex as fixed effects. Likewise, to model post-fledging activity, we fitted an LMM with post-fledging ODBA (random intercept of the ODBA model described above) as response variable and NDVI, sex and ODBA in the nest as fixed effects. The fledging age was modelled by fitting an LMM with nestling ODBA, nest exposition, body condition and sex as fixed effects. Foray patterns were assessed by fitting a Poisson GLMM for the number of forays and LMMs each for mean foray duration (square root-transformed) and mean maximum foray distance (log-transformed) as response variables with post-fledging ODBA, NDVI and sex as fixed effects. Finally, to assess factors influencing the individual age at departure, we fitted an LMM with post-fledging ODBA, NDVI, age at fledging and sex as fixed effects. To account for the non-independence of data between nestlings from the same brood or territory, as well as across years, we included territory, individual ID and year as crossed random effects in all models.

Explanatory variables were checked for collinearity using pairplots and Pearson’s correlation coefficient r. Because none of the explanatory variables showed a correlation of r ≥ 0.7, all were retained in the models. In addition, all continuous explanatory variables were centred and scaled. Models were checked for convergence of the Markov chains with Brooks-Rubin-Gelman diagnostics [45] and for validity using posterior predictive checks (package ‘shinystan’; [46]). Spatial autocorrelation in the residuals was checked using Moran’s I (package ‘*ape*’; [47]) and bubbleplots (package ‘*sp*’, [48]) and no autocorrelation was detected. Models were created based on ecological hypotheses and no model selection was performed. Inference and predictions were based on simulated Bayesian posterior distributions using 10’000 simulations (4 chains of 5’000 iterations with a burn-in of 2’500 each). Additionally, posterior probabilities (PP) were calculated for all model estimates, indicating the probability of the estimates to differ from zero. PP ≥ 0.95 indicated strong support for an existing effect, while 0.95 > PP > 0.9 indicated a clear tendency for an existing effect. We assume that the tendencies represent existing effects which show higher uncertainty due to the limited sample size in this study. Therefore, we report tendencies as slight effects in the results section. Unless stated otherwise, model parameters and derived parameters are given as posterior means with 95% credible intervals (95% CrI).

## 4. Results

The golden eagle nestlings weighed 3.52 ± 0.53 sd kg (range: 2.21 to 4.60 kg; N = 35) at tagging. Females weighed 3.87 ± 0.46 sd kg (range: 2.88 to 4.60 kg; N = 17) and males weighed 3.20 ± 0.37 sd kg (range: 2.21 to 3.70 kg; N = 19). Nestling ODBA (daily mean of all 7.9 sec bursts) in the 11 days prior to fledging was 0.05 ± 0.01 sd *g* (gravitational force; range: 0.03 to 0.08 *g*; N = 35) and increased to post-fledging ODBA of 0.08 ± 0.01 sd *g* (range: 0.06 to 0.10 *g*; N = 32) in the first 45 days after fledging. Nestlings fledged at an age of 77.9 ± 6.2 sd days (range: 62 to 90 days; N = 35). All individuals undertook multiple forays (mean = 11.9 ± 6.7 sd forays; range 3 to 29; N = 32) during the post-fledging period before departing, lasting for 1.3 ± 1.0 sd days (range: 0.1 to 4.2 days; N = 32) directing them to areas in 22.4 ± 7.4 sd km (range: 13.3 to 137.2 km; N = 32) distance from the parental territory. No juvenile died between fledging and departure from the parental territory. Independence and thus, final departure from the parental territory to dispersal occurred at an age of 280.6 ± 40.6 sd days (range: 185 to 328 days; N = 32).

### Nestling period and fledging age

Nestling ODBA was not associated with the two environmental variables of the nest surrounding, nest exposition and NDVI (Table 1). Instead, male nestlings showed higher ODBA than female nestlings. Accounting for environmental variables and sex, the model showed slight support for an effect of nestling condition on nestling ODBA. ODBA was higher in nestlings in good body condition compared to nestlings in poorer body condition (Table 1, Figure 1a).

**Table 1.** Estimates of the Linear Mixed Models investigating factors affecting nestling ODBA (11 days pre-fledging) and post-fledging ODBA (45 days post-fledging). Means (Estimates), 95% CrI and Bayesian posterior probabilities (PP) are given. Fixed effects with PP ≥ 0.95 are printed in bold; with 0.95 > PP > 0.9 in bold and italics. Nestling ODBA: n= 35 individuals from 27 territories. Post-fledging ODBA: n = 32 individuals from 26 territories. Standard deviations and 95% CrI of random effects are shown in Table S2.

| Model | Fixed Effects | Estimates | 95% CrI | PP (< or > 0) |
| --- | --- | --- | --- | --- |
| Nestling ODBA | Intercept | **0.052** | **0.041 to 0.061** | **1.000** |
|  | Nest exposure (south) | -0.001 | -0.004 to 0.001 | 0.888 |
|  | NDVI | < 0.000 | -0.002 to 0.003 | 0.530 |
|  | Sex (male) | **0.003** | **-0.001 to 0.008** | **0.958** |
|  | Body condition | **0.001** | **-0.001 to 0.004** | **0.913** |
| Post-fledging ODBA | Intercept | **0.077** | **0.067 to 0.085** | **1.000** |
|  | Nestling ODBA | **0.005** | **-0.001 to 0.010** | **0.935** |
|  | NDVI | 0.001 | -0.002 to 0.005 | 0.752 |
|  | Sex (male) | **0.005** | **-0.001 to 0.011** | **0.955** |

**Figure 1.**
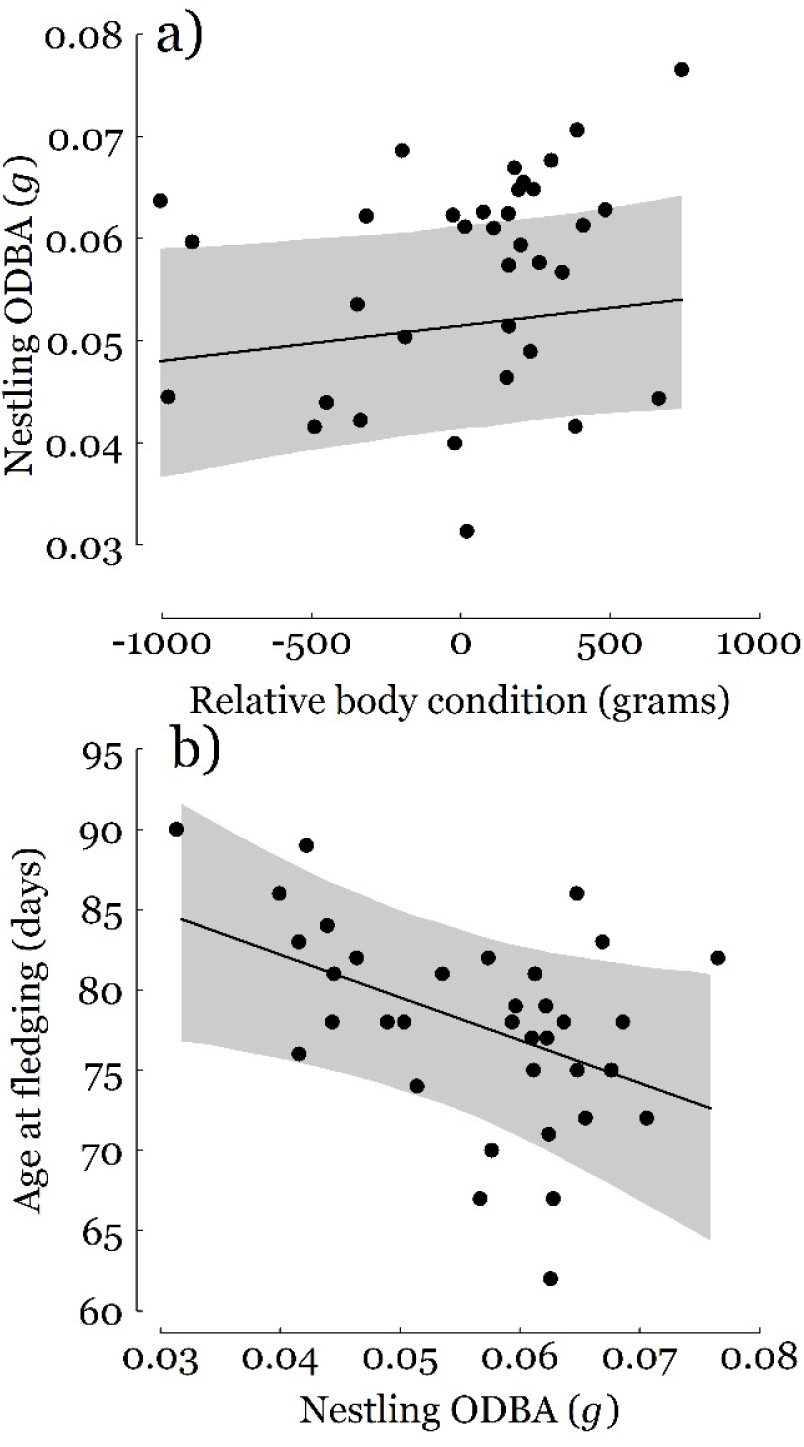
**a)** Nestling activity (average over 11 days pre-fledging; measured as ODBA in gravitational force *g*) of golden eagles in relation to relative nestling body condition (residuals from mean body weight of the respective sex in grams) and **b)** Age at fledging (in days from hatching) of golden eagle nestlings in relation to nestling activity (average over 11 days pre-fledging; measured as ODBA in gravitational force *g*). The solid line shows the mean and shaded areas 95% CrI of model predictions for females with all other model predictors set to their mean values. Filled circles show raw data points for both sexes. N = 35 individuals from 27 territories and four years.

Fledging age was strongly related to nestling ODBA and nest exposition. Nestlings with high ODBA fledged earlier than individuals with low ODBA (Table 2, Figure 1b) and individuals from southern exposed nests fledged earlier than individuals from northern exposed nests (Table 2). When accounting for the effect of nestling ODBA, nestling body condition did not show an effect on fledging age. Yet, male nestlings tended to fledge earlier than females in addition to their earlier fledging due to the increased ODBA (Table 2).

**Table 2.** Estimates of the Linear Mixed Models investigating factors affecting age at fledging, exploratory behaviour (number of forays, Generalized Linear Mixed Model); foray duration (sqrt-transformed), foray distance (log-transformed)), and age at departure. Means (Estimates), 95% CrI and Bayesian posterior probabilities (PP) are given. Fixed effects with PP ≥ 0.95 are printed in bold; with 0.95 > PP > 0.9 in bold and italics. Age at fledging: n= 35 individuals from 27 territories; Forays and age at departure: n = 32 individuals from 26 territories. Standard deviations and 95% CrI of random effects are shown in Table S2.

| Model | Fixed effects | Estimates | 95% CrI | PP |
| --- | --- | --- | --- | --- |
| Age at fledging | Intercept | **78.433** | **72.932 to 84.129** | **1.000** |
|  | Nestling ODBA | **-2.673** | **-5.123 to -0.196** | **0.984** |
|  | Nest exposure (south) | **-2.187** | **-3.697 to -0.636** | **0.995** |
|  | NDVI | 0.029 | -1.545 to 1.684 | 0.507 |
|  | Sex (male) | **-2.223** | **-5.391 to 0.787** | **0.928** |
|  | Body condition | -0.817 | -2.413 to 0.705 | 0.854 |
| Number of forays | Intercept | **2.592** | **1.996 to 3.239** | **1.000** |
|  | Post-fledging ODBA | **0.191** | **-0.090 to 0.471** | **0.919** |
|  | NDVI | -0.009 | -0.251 to 0.226 | 0.529 |
|  | Sex (male) | **-0.324** | **-0.736 to 0.070** | **0.944** |
| Mean foray duration | Intercept | **1.271** | **0.987 to 1.587** | **1.000** |
|  | Post-fledging ODBA | -0.014 | -0.159 to 0.189 | 0.561 |
|  | NDVI | -0.012 | -0.150 to 0.176 | 0.561 |
|  | Sex (male) | **-0.342** | **-0.645 to -0.059** | **0.990** |
| Mean foray distance | Intercept | **3.228** | **3.030 to 3.434** | **1.000** |
|  | Post-fledging ODBA | -0.016 | -0.138 to 0.102 | 0.606 |
|  | NDVI | -0.006 | -0.120 to 0.106 | 0.538 |
|  | Sex (male) | **-0.313** | **-0.530 to -0.098** | **0.998** |
| Age at departure | Intercept | **283.629** | **257.674 to 311.178** | **1.000** |
|  | Post-fledging ODBA | **-10.787** | **-27.874 to 6.191** | **0.901** |
|  | NDVI | -3.180 | -20.087 to 14.143 | 0.656 |
|  | Sex (male) | -5.923 | -32.433 to 20.087 | 0.687 |
|  | Age at fledging | **-12.150** | **-28.303 to 4.720** | **0.926** |

### Post-fledging period and departure

Females tended to undertake more forays after fledging than males (Table 2). Females also spent more time and covered longer distances on individual forays than males (Table 2). NDVI did neither affect foray behaviour nor age at departure (Table 2). All individuals increased ODBA over the first 45 days post-fledging (Table S1; Figure S1). As in the nest, males also showed higher post-fledging ODBA than females (Table 1). The model showed slight support for fledglings with high nestling ODBA also showing high post-fledging ODBA (Table 1, Figure 2) and high post-fledging ODBA was slightly associated with an increased number of post-fledging forays, but not with the duration and distance of the forays (Table 2, Figure 3a). Fledglings with high post-fledging ODBA showed a slight effect to depart at a younger age than fledglings with a low post-fledging ODBA (Table 2; Figure 3b). When accounting for this ODBA effect, juveniles that fledged later than their peers tended to depart after a shorter post-fledging period than their peers (Table 2). This can be explained by the fact that individuals fledging later also fledged later in the season (r = 0.51; GLMM: Estimate = 0.30; 95% CrI = 0.09 to 0.52), leaving them on average less time until emigration. When propagating the effect of nestling activity to the age of emigration, nestlings with the highest activity emigrated at a 16 days younger age than the nestlings with the lowest activity. Even though sexes differed in fledging age and ODBA patterns both in the nestling and in the post-fledging period, males and females did not differ in age at departure to dispersal (Table 2).

**Figure 2.**
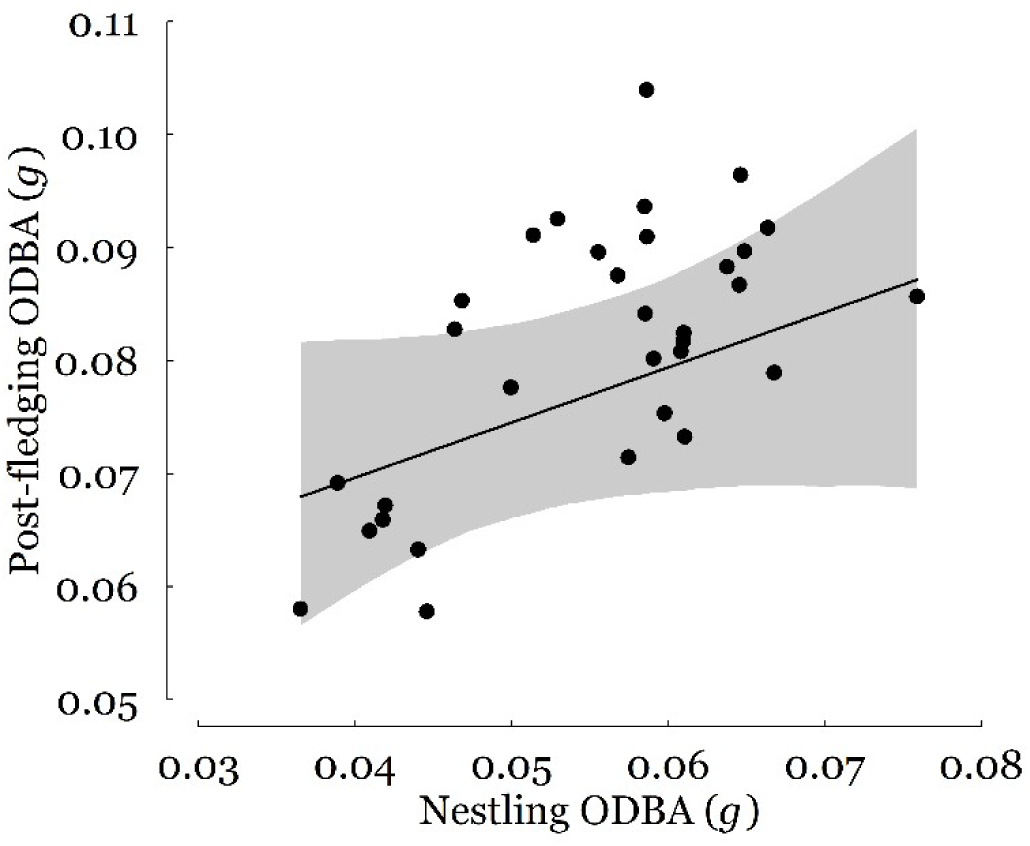
Post-fledging activity (over 45 days post-fledging; measured as ODBA in gravitational force *g*) of juvenile golden eagles in relation to their nestling activity (over 11 days pre-fledging; measured as ODBA in gravitational force *g*). The solid line shows the mean and the shaded area 95% CrI of model predictions for females with all other model predictors set to their mean values. Filled circles represent raw data points for both sexes. N = 32 individuals from 26 territories and four years.

**Figure 3.**
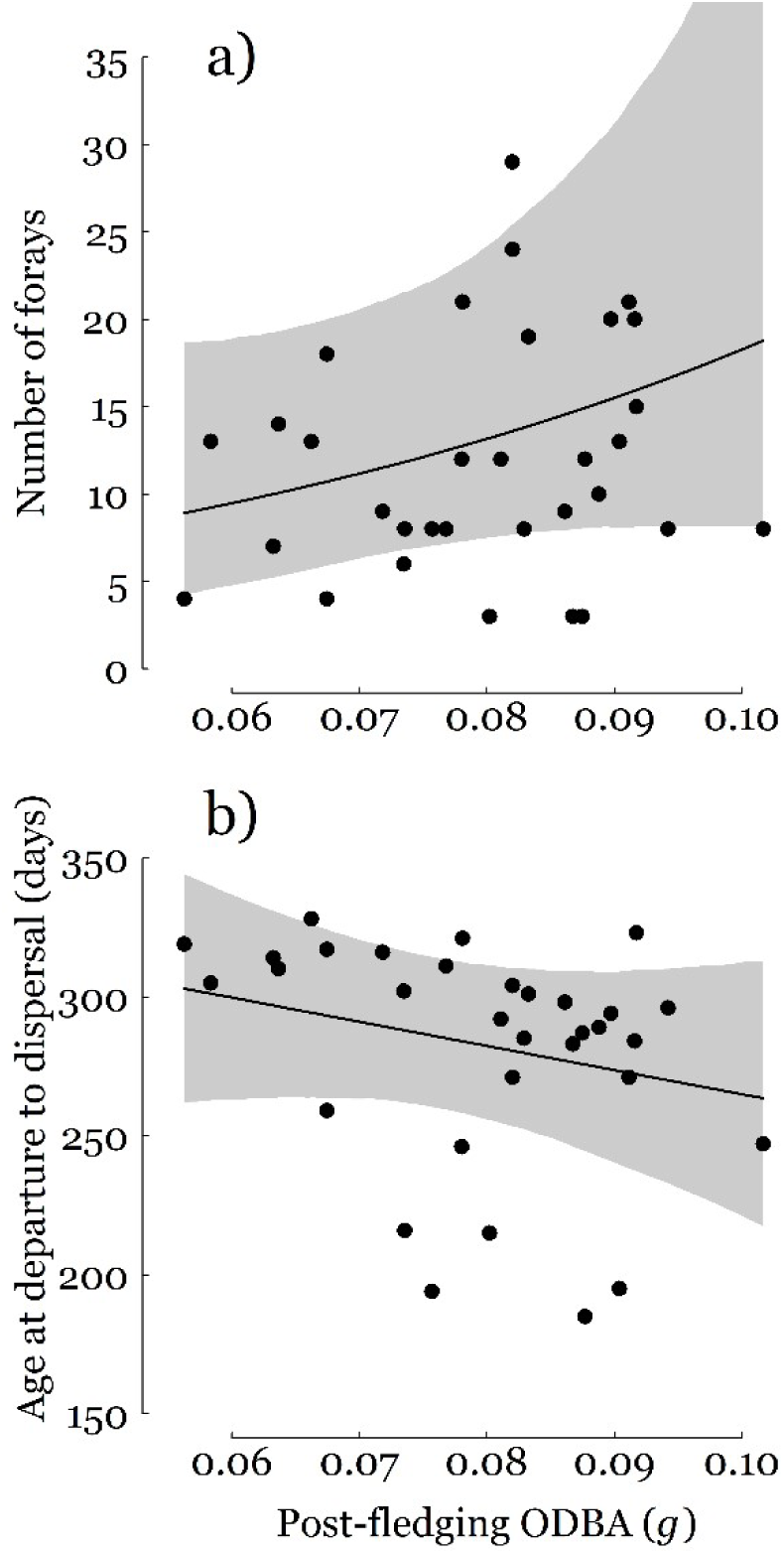
**a)** Number of forays and **b)** age at departure to dispersal of juvenile golden eagles in relation to post-fledging activity (over 45 days post-fledging; measured as ODBA in gravitational force *g*). The solid line shows the mean and the shaded area 95% CrI of model predictions for females with all other model predictors set to their mean values. Filled circles show raw data points for both sexes. N = 32 individuals from 26 territories and four years.

## 5. Discussion

Our results, despite some reduced certainty, provide novel insights into the correlational chain from conditions experienced in the nestling period over the first life-history transition of fledging to the post-fledging period and further to the timing of natal dispersal. Nestling golden eagles in good body condition tended to be more active than nestlings in poorer condition, and high activity was associated with fledging at a young age. Active nestlings continued to be more active during the post-fledging period, undertaking more forays and ultimately departing earlier for natal dispersal than passive nestlings. Thus, the results show evidence for an association between nestling condition giving rise to a nestlings’ activity phenotype which seemed to persist throughout the post-fledging period to affect departure timing.

### Origin of the activity phenotype

Individual activity, measured as overall dynamic body acceleration (ODBA), is strongly driven by the frequency of high-motion behaviours [26]. While behaviours such as standing or roosting show activity levels close to basal metabolic rate at rest, behaviours associated with movement such as flight result in largely increased activity levels [21,49]. In the nest, the set of behaviours include mainly low energy behaviours, but nestlings increasingly walk, spread and flap wings when approaching the time of fledging [50]. Likewise, during the first weeks of the post-fledging period, birds allocate more and more time to flight at the expense of roosting [21,22]. Birds with consistently higher activity levels were thus likely to show more behaviours associated with locomotion, particularly flight, both in the nest and in the early post-fledging period.

The individual differences in activity levels, i.e. the activity phenotype expressed in the nest and in the post-fledging period, can either arise from environmental conditions experienced in the nest [51] or from intrinsic differences in the development of behaviours [52] and tendencies to react to stimuli [53]. The slight correlation of activity levels with body condition found in this study suggests that the activity phenotypes developed in the nest were at least partly driven by environmental conditions. Yet, a large part of the variation in nestling activity remained unexplained, suggesting that physiological or developmental characteristics were not directly related to the environment nor contributed to the observed behavioural phenotypes. The observed consistent sex-differences in activity levels, timing of fledging and exploration behaviour provide support for intrinsic differences in behavioural phenotypes. Based on a generally positive relationship of ODBA with body mass [26], we would have expected males, the smaller sex, to show lower activity levels than females. The inverse relationship observed indicates that there were intrinsic differences in activity between sexes or that females needed more time to physiologically develop to their larger size and to reach the same level of high-activity behaviour as males did [23,54]. Arguably, the different size of sexes might also affect activity measurements, for example by different effects of turbulences on the tags when flying.

The lack of effect of biomass productivity (NDVI) on activity levels, fledging age or movement characteristics at any stage investigated could be interpreted as support for intrinsic behavioural differences. However, it is possible that NDVI represents a poor proxy for environmental quality of natal territories in golden eagles. It is conceivable that golden eagle prey abundance is not strongly linked to NDVI. This is underpinned by the lack of relationship between NDVI and nestling body condition (LMM: Estimate: -84.5, 95% CrI: -245.2 to 79.5). In addition, other environmental factors such as the topographic configuration of territories and associated flight conditions may be more decisive for food accessibility (parental provisioning), juvenile flight propensity and locomotory development [31,40].

The effect of nest exposure on the timing of fledging corroborates the importance of topography and thermal conditions for behavioural development. Early fledging observed in southern exposed nests could indicate that southern slopes offer improved flight conditions to parents for provisioning but also to juveniles allowing them to fledge earlier. It also implies that optimal nest sites are limiting or thermal conditions need to be traded against other factors affecting nest site selection. While these results suggest combined intrinsic and environmental effects driving flight activity and development, the direct and indirect contribution of specific factors deserves further investigation.

### Carry-over of the activity phenotype

There is ample evidence for a strong link between nestling condition and post-fledging survival in a large range of species [3,16]. Our findings that golden eagle nestlings with high activity levels fledged at a young age and tended to show high post-fledging activity levels suggests that this effect is rooted in behavioural differences. In long-lived species with high post-fledging survival rates such as the golden eagle [55], nestling activity phenotypes seem to carry-over to affect the behaviour throughout the entire post-fledging stage. In other species, carry-over effects of nestling condition on behaviour and fitness correlates later in life were often attributed to physiological conditions such as oxidative stress, corticosterone levels or telomere length [10,11]. We now show that nestling condition can also be reflected by the nestlings’ activity phenotype and that the behavioural differences underlying the activity phenotype can carry-over to later life stages.

Forays outside the parental territory may be a beneficial move to attenuate the risks of dispersal: Forays allow individuals to obtain information on the surrounding environment and gain experience before leaving the parental territory permanently [19]. During forays, juveniles may assess the relative habitat quality of their natal range [56], practice to localise suitable feeding grounds and beneficial flight conditions [57], or learn to avoid agonistic behaviours of conspecific and anthropogenic threats once departed [58]. More active juveniles may thus likely gain fitness benefits from undertaking more forays if the knowledge gain carries over to the dispersal phase.

The subsequent permanent departure from the natal area is a fundamental changepoint in the life of juvenile animals, as it entails leaving the known environment, the ending of parental care and concomitant shelter from predation or conspecific aggression [19]. This life-history transition to natal dispersal is thus risky and should only occur at a point when skills are developed to a degree that allow to deal with these risks [20]. Juveniles may thus have an interest in staying with their parents for as long as possible to gain experience, reach behavioural maturity and a good nutritional status [22]. However, pampering their offspring might cause costs to parents that can be reduced by an early emigration of juveniles [59]. In addition, a timely emigration possibly also benefits juveniles, as it can support the detection and monopolisation of patchy food resources, the acquisition of high-quality floater areas or higher competitive ability in the social system [19,60]. The fact that individuals with high activity levels emigrated early suggests that they reach this point earlier than individuals with low activity levels. This is supported by a strong positive correlation between overall post-fledging activity levels and the increase of activity levels (correlation between random intercept and random slope of ODBA: Pearson’s r = 0.89). The accumulated differences associated with the high activity phenotype at departure may thus lower dispersal costs to increase survival [9,20], allow to overcome energetic constraints of dispersal [25,61], and facilitate an optimal dispersal outcome by increasing the likelihood of finding and settling in high-quality habitats early in life [8]. The carry-over effect of activity differences from the nest to the post-fledging period and its subsequent translation to the start into the next life stage may ultimately have pursuing effects on post-dispersal stages [8,14].

## 6. Conclusions

Even with our low sample sizes, we found evidence for carry-over effects translating behaviours associated with early-life conditions over two life-history transitions. The well-known effect of early-life food conditions on nestling body condition [11,62] was associated with an activity phenotype still present in the subsequent post-fledging life stage and which carried over to affect timing of life-history transitions and movement decisions of juvenile birds. When propagating differences in nestling activity up to emigration, nestlings with the highest activity emigrated 16 days earlier compared to nestlings with the lowest activity. Our study therefore adds to the growing literature showing that behavioural phenotypes shaped by early-life conditions can persist and thereby represent a key mechanism linking early-life conditions with future survival and reproduction. However, further research shedding light on the phenotypic variation in specific behaviours underlying the activity phenotype will refine our understanding of the behavioural mechanisms linking early-life conditions to individual fitness in long-lived animals and allow for better predictions of their population-level consequences.

## Supporting information

Electronic supporting information

## Acknowledgments

We thank the Department of Wildlife and Fishery Service Grisons, Switzerland, the mountain rescue teams of the Guardia di Finanza, Italy, the forestry and game wardens of South Tyrol/Alto Adige, Italy, the Stelvio National Park, the Adamello Regional Park (Anna Bonettini), the Paneveggio Pale di San Martino Natural Park (Gilberto Volcan and Piergiovanni Partel), the Sondrio Province (Maria Ferloni), ISPRA (Lorenzo Serra) and the Regione Lombardia (Guido Pinoli) for their support for monitoring, ringing and tagging the birds. In specific, we want to mention Claudio Angeli, Geni Ballat, Klaus Bliem, Daniel Bundi, Adriano Greco, Hannes Jenny, Ueli Jörimann, Alessandro Mercogliano, Carlo Micheli, Stefan Rauch, Andrea Roverselli, Romano Salis. Then, we are grateful to Franziska Lörcher and Daniel Hegglin for their help with constructing harnesses, Fränzi Korner for statistical advice, Jérôme Guélat for GIS support, and Marta Burri and Juanita Olano Marin for molecular sex identification of feather samples. Furthermore, we thank Ilse Storch and Michael Schaub for their comments on an earlier version of this manuscript. This study would not have been possible without Martin Wikelski who provided the tags.

## Ethical statement

Handling, tagging and marking of golden eagle nestlings in Switzerland was carried out under the permit by the Federal Food Safety and Veterinary Office Grisons (licence No. GR 2017_06, GR 2018_05E, GR 2019_03E). In Italy, the permissions for handling, tagging and marking were obtained from autonomous region of South Tyrol (Dekret 12257/2018 and Dekret 8788/2020), as well as from the Regione Lombardia and Sondrio Province for ringing and tagging in Lombardia and South Tyrol by ISPRA (Istituto Superiore per la Protezione e la Ricerca Ambientale) with the Richiesta di autorizzazione alla cattura di fauna selvatica per scopi scientifici (l.r. 26/93). All procedures followed the ASAB/ABS guidelines for the ethical treatment of animals in behavioural research and teaching and all applicable international, national, and/or institutional guidelines for the care and use of animals were followed. The handling of birds was performed with maximum care and disturbance to nests kept to a minimum. Ethical approval for involving animals in this study was received through the application procedure for ringing permits and the scientific commission of the Swiss Ornithological Institute.

## Data accessibility

Data and code are available on the vogelwarte.ch Open Repository and Archive: https://doi.org/10.5281/zenodo.15576191.

## Authors’ contributions

S.-S.Z.: Conceptualization, Data curation, Formal analysis, Investigation, Methodology, Visualization, Writing – original draft. M.U.G.: Conceptualization, Funding acquisition, Investigation, Methodology, Project administration, Resources, Supervision, Validation, Writing – review & editing. J.S.H.: Conceptualization, Data curation, Investigation, Methodology, Project administration, Resources, Supervision, Validation, Writing – review & editing. E.B.: Data curation, Funding acquisition, Project administration, Resources, Writing – review & editing. K.S.: Conceptualization, Data curation, Funding acquisition, Project administration, Resources, Validation, Writing – review & editing. W.F.: Data curation, Funding acquisition, Project administration, Resources, Writing – review & editing. D.J.: Data curation, Funding acquisition, Project administration, Resources, Validation, Writing – review & editing. M.T.: Conceptualization, Formal analysis, Funding acquisition, Investigation, Methodology, Project administration, Resources, Supervision, Validation, Visualization, Writing – review & editing.

