## Supplementary material for "Cascading carry-over effects of early activity phenotypes in golden eagles": Electronic supporting information

**Electronic Supplementary Material**

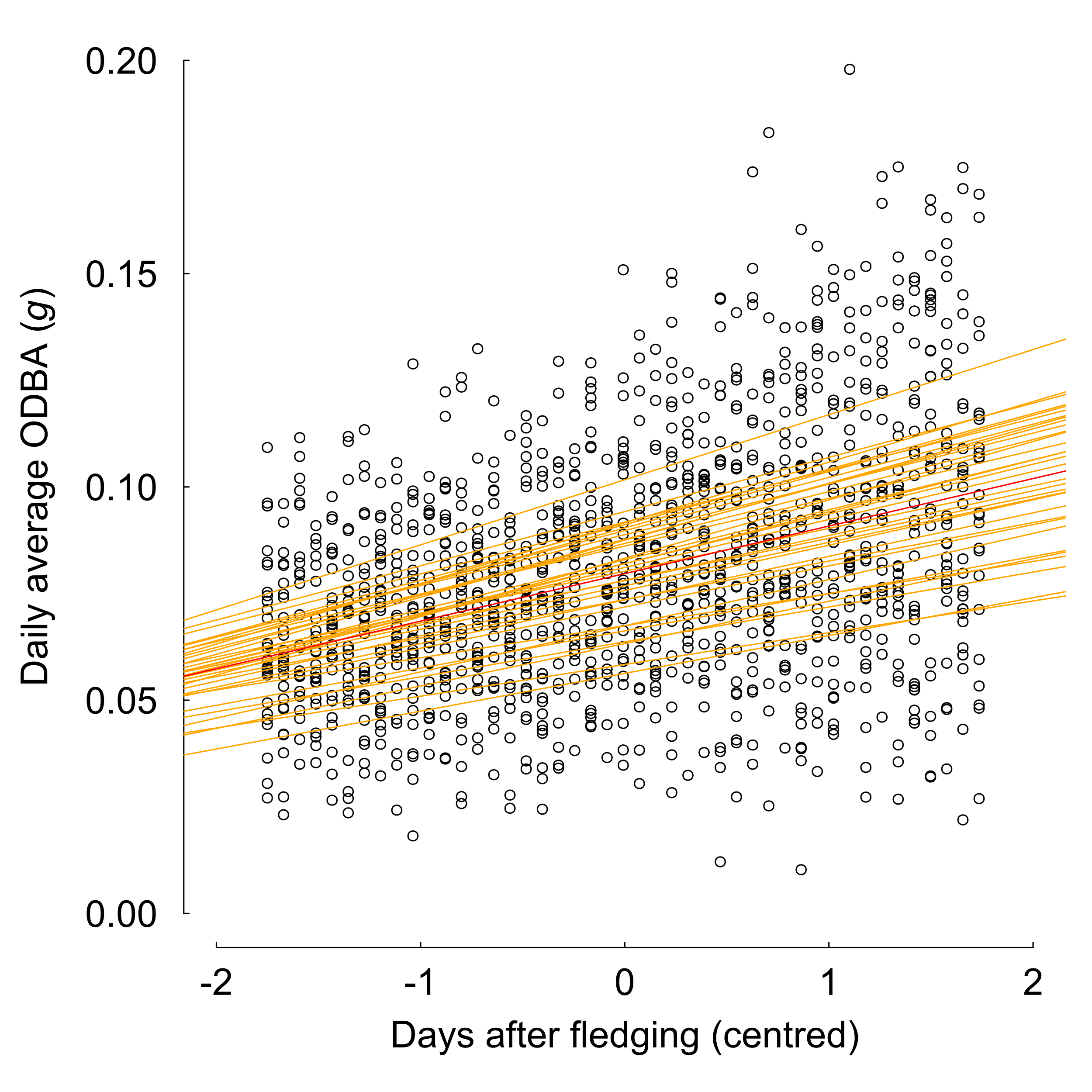

**Figure S1.** Development of individual daily activity (ODBA in gravitational force *g*) for all 32 individuals during the 45 days after fledging. Orange lines show individual regression lines and the red line the population mean of the standardised random intercepts and slopes of a linear mixed-effects model modelling individual ODBA in response to day after fledging and sex (Table S1).

**Table S1.** Estimates of the Linear Mixed Models assessing the development of daily activity (ODBA in gravitational force *g*) in the 11 days before fledging (Individual daily nestling ODBA) and the 45 days after fledging (Individual daily post-fledging ODBA). Means (Estimates), 95% CrI and Bayesian posterior probabilities (PP) of fixed effects, as well as standard deviations (sd) and 95% Credible Intervals of random effects are given. Fixed effects with PP ≥ 0.95 are printed in bold; with 0.95 > PP > 0.9 in bold and italics. Individual daily nestling ODBA: n= 35 individuals; Individual daily post-fledging ODBA: n = 32 individuals.

| **Model** | **Fixed Effects** | **Estimates** | **95% CrI** | **PP (< or > 0)** |
| --- | --- | --- | --- | --- |
| Individual daily nestling ODBA | Intercept | **0.054** | **0.049 to 0.060** | **1.000** |
|  | Day | 0.0002 | -0.002 to 0.002 | 0.600 |
|  | Sex (male) | 0.004 | -0.003 to 0.012 | 0.860 |
|  | sd Individual:day (Intercept) | -1.541e^-5^ | -4.214e^-5^ to 6.172e^-6^ |  |
|  | sd Individual:day (Slope) | 1.962e^-5^ | 6.075e^-6^ to 4.168e^-5^ |  |
| Individual daily post-fledging ODBA | Intercept | **0.080** | **0.074 to 0.086** | **1.000** |
|  | Day | **0.011** | **0.010 to 0.013** | **1.000** |
|  | Sex (male) | 0.002 | -0.006 to 0.011 | 0.719 |
|  | sd Individual:day (Intercept) | 2.101e^-5^ | 1.184e^-6^ to 4.791e^-5^ |  |
|  | sd Individual:day (Slope) | 8.805e^-6^ | 1.431e^-6^ to 2.267e^-5^ |  |

**Table S2.** Standard deviations (sd) and 95% Credible Intervals of random effects for all models.

| **Model** | **Random Effects** | **SD** | **95% CrI** |
| --- | --- | --- | --- |
| Nestling ODBA | Year | 9.22e^-5^ | 1.81e^-5^ to 3.12e^-4^ |
|  | Territory ID | 1.17e^-5^ | 1.39e^-8^ to 4.32e^-5^ |
| Post-fledging ODBA | Year | 5.36e^-5^ | 1.38e^-8^ to 2.58e^-4^ |
|  | Territory ID | 3.23e^-5^ | 1.45e^-7^ to 8.75e^-5^ |
| Age at fledging | Year | 26.9 | 3.33 to 103.11 |
|  | Territory ID | 2.46 | 0.00 to 12.14 |
| Number of forays | Year | 0.26 | 0.00 to 1.53 |
|  | Territory ID | 0.26 | 0.10 to 0.54 |
| Mean foray duration | Year | 0.038 | 0.00 to 0.24 |
|  | Territory ID | 0.08 | 0.00 to 0.21 |
| Mean foray distance | Year | 0.01 | 0.00 to 0.10 |
|  | Territory ID | 0.02 | 0.00 to 0.07 |
| Age at departure | Year | 250.62 | 0.08 to 1667.76 |
|  | Territory ID | 1211.08 | 92.11 to 2537.75 |
